# AIMP1-derived peptide secreted from hair follicle stem cells activates dermal papilla cells to promote hair growth

**DOI:** 10.1101/2022.02.24.481761

**Authors:** YounHa Kim, Ho Lee, Doyeun Kim, Soon Sun Bak, Ina Yoon, Ralf Paus, Seongmin Cho, Seung Jae Jeong, Yoon Jeon, Min Chul Park, Ji Won Oh, Jung Min Park, Sang Bum Kim, Young Kwan Sung, Sunghoon Kim

## Abstract

Hair follicle stem cells (HFSCs) and dermal papilla cells (DPCs) are crucial in the biogenesis and maintenance of hair follicles (HFs). In this study, a fragment derived from aminoacyl-tRNA synthetase-interacting multifunctional protein1 (AIMP1) was secreted from HFSCs to activate DPCs to maintain hair follicle homeostasis. A histological analysis revealed that AIMP1 levels in hair follicles decreased with hair loss. Hair regrowth in AIMP1-induced mice was faster than that in non-induced mice. Deletion mapping revealed 41 amino acids (TN41, aa 6-46) as the active region of AIMP1. The N-terminal peptide fragment of AIMP1 generated by MMP1 was secreted from Wnt-treated HFSCs to activate DPCs via FGFR2. TN41 activated Akt and ERK, increased β-catenin, and enhanced DPCs activation. TN41 also promoted hair shaft elongation in cultured human hair follicles and improved the hair-inducing activity of cultured DPC spheroids. In summation, the AIMP1 fragment secreted from HFSCs appears to stimulate active hair regrowth through activating DPCs.

## INTORDUCTION

HFs undergo cyclical transformations and progress through the stages of growth (anagen), regression (catagen), and rest (telogen). During the anagen phase, HFs produce hair shafts. HFs reset and prepare HFSCs and DPCs during the catagen and telogen phases. HFSCs and DPCs then communicate via signaling molecules such as Wnt7b, FGF7, and Noggin to initiate the next growth phase and create a new hair shaft for hair homeostasis ^1^.

HFSCs, which reside in the bulge area of the HF, sustain cyclic hair regrowth over repeated cycles ^2^. Communication between HFSCs and their niches is reciprocal because stem cells regulate homeostasis and maintenance of niches ^3^. Mouse HFSCs generally do not display apparent decreases in numbers ^4^; however, aging exhibits imbalanced cytokine signaling with their niches and diminished colony-forming capability ^5^. Loss of communication between HFSCs and their niches decreases hair density and increases hair thinning in mammals that live relatively long period ^6^.

DPCs are widely recognized as the center that triggers hair cycling via a paracrine signaling mechanism ^7^. Specifically, crosstalk between DPCs and HFSCs is essential for ensuring postnatal hair growth and HF cycling ^8^. DPCs are physically closest to HFSCs during telogen, and various factors secreted by these cells are likely involved in the progression from telogen to anagen. Understanding the mechanisms of this telogen-to-anagen transition and anagen maintenance is key to ensuring successful hair loss treatment.

Previous studies have shown that fibroblast growth factor (FGF) receptor (FGFR) signaling plays an important role in regulating the hair cycle. *In situ* hybridization was used to assess the expression of all four FGFRs during the growth phase. Among them, FGFR1 and FGFR2 are abundantly expressed in DPCs throughout the anagen phase, and FGFR2-mediated FGF signaling modulates the hair cycle ^9^.

AIMP1 was originally identified as a member of the mammalian multi-tRNA synthetase complex ^10^. AIMP1 is secreted in response to cytokine stimulation and exerts distinct extracellular activities depending on the target cells ^11, 12, 13, 14, 15^. Human AIMP1 mutations are associated with severe neurodegenerative disorders. Mutations in AIMP1 appear related to the development and maintenance of axon cytoskeleton integrity and regulation of neurofilaments ^16, 17^. Furthermore, AIMP1 is associated with cellular homeostasis. Macrophages secrete AIMP1 following stimulation with tumor necrosis factor (TNF)-α to enhance wound healing. This process is mediated by AIMP1-induced fibroblast proliferation and collagen synthesis via Akt and ERK activation ^18^. Deletion-mapping analysis revealed that the N-terminal domain of AIMP1 stimulates fibroblast and mesenchymal stem cell (MSC) proliferation ^19, 20^. FGFR2 on MSCs is a receptor for the N-terminal fragment of AIMP1. Additionally, the expression of AIMP1 is high in DPCs and dermal sheath, suggesting that AIMP1 could play a new role in the skin or HFs ^21^.

## RESULTS

### AIMP1 protects against hair loss by inducing anagen

Aged mice exhibited diffuse and symmetric patterns of hair loss on their backs. These patterns typically became obvious in the most dorsal area, where an uneven or linear hair loss pattern formed and spread toward their flanks. Similarly, hair loss in C57BL/6 mice was apparent approximately 16 months after birth, and we used 16-month-old mice for histological analysis of HFs in the sparse hair (SH) and normal (H) regions (Figure 1a, red and yellow circles, respectively). As expected, skin in the SH region exhibited a significant decrease in HFs (Figure 1a). To determine the functional relevance of AIMP1 in HF maintenance, we compared the expression levels of AIMP1 between normal and SH regions using immunofluorescence (IF) staining. AIMP1 was originally distributed evenly in all tissues; however, AIMP1 expression was especially enriched in the bulge at telogen phase (Figure 1b). AIMP1 levels decreased in the bulges isolated from the SH region (Figure 1b and Figure S1a). The fluorescence staining intensity (FI) of AIMP1 in the H regions was three-fold higher than that in the SH region (Figure 1b). As a negative control, the expression level of glutamyl-prolyl-tRNA synthetase (EPRS), a component of the multi-tRNA synthetase complex, together with AIMP1, was similar between the two regions (Figure S1b) ^22^. HFSCs reside in the bulge, and AIMP1 is an enriched bulge region. Therefore, we examined the levels of HFSC markers, including SOX9 and K15, along with AIMP1 using immunohistochemistry. Interestingly, AIMP1 colocalized with these markers in bulge cells (Figure S1c).

**Figure 1.**
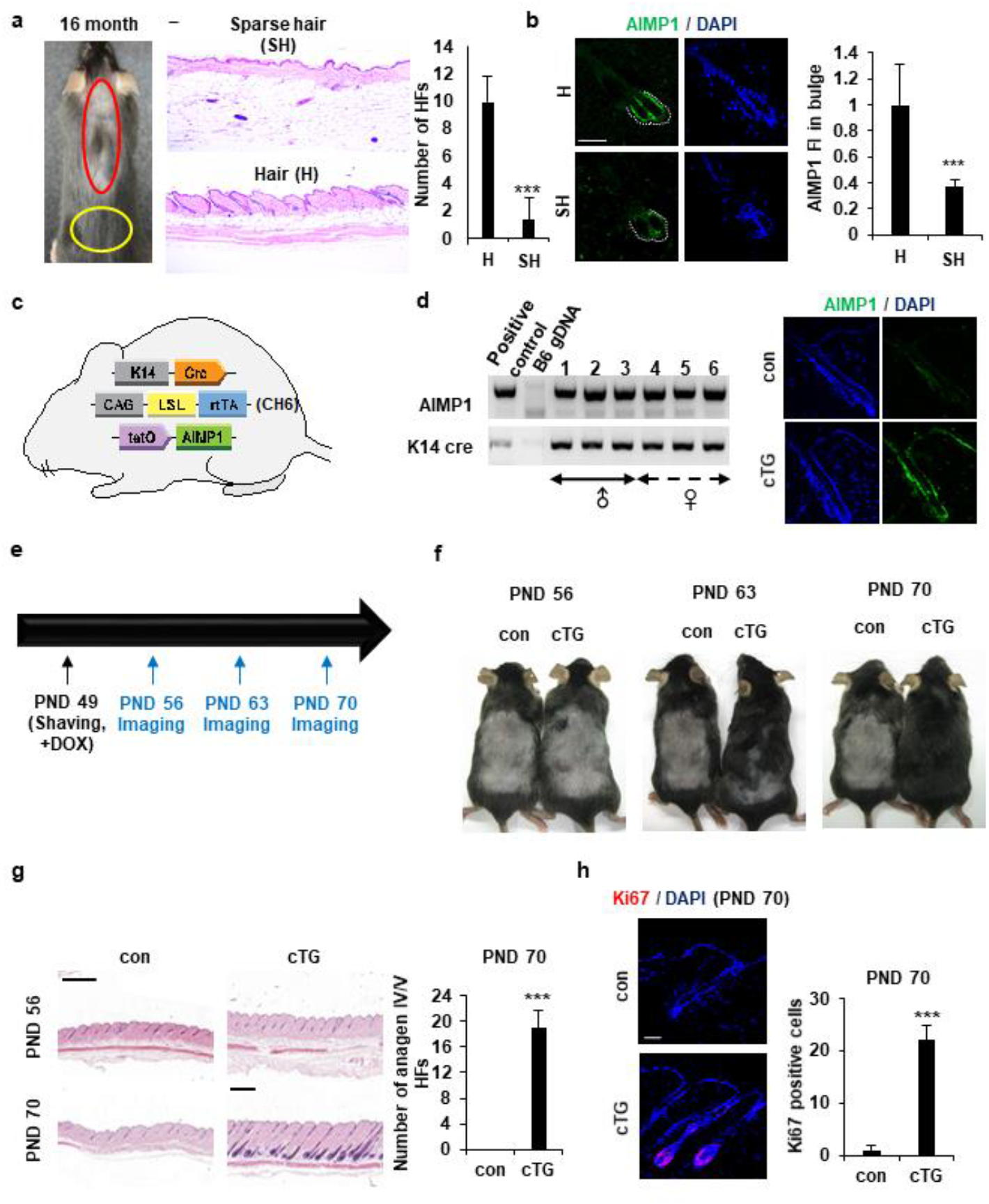
AIMP1 for HF maintenance. (**a**) Representative and histological pictures of C57BL/6 (n = 4). HFs counts from each region. (**b**) IF images of AIMP1. Relative FI was measured using Image J. (**c**) A schematic diagram of the skin-specific AIMP1 inducible mouse. (**d**) Validation of cTG by genotyping and IF staining of AIMP1. (**e**) The design for analyzing hair growth. (**f**) Representative pictures of mice taken at indicated days (n = 2). (**g**) H&E image from each mouse. HFs counts from each mouse. (**h**) IF staining of Ki67 and counts of Ki67 positive cells from each mouse. The bars: 50 μm (**a, b and h**) and 500 μm (**g**). Error bars: mean ± SD. ***: P < 0.001

To determine whether AIMP1 influences HF maintenance, we generated KRT14cre-AIMP1 conditional transgenic (cTG) mice in which AIMP1 expression was induced in a skin-specific manner (Figure 1c). Generation of AIMP1 cTG mice was verified by using genotyping and IF staining (Figure 1d). Mice were clipped on postnatal day (PND) 49 when the second telogen normally takes place, and a picture was taken every week thereafter (Figure 1e) ^23^. AIMP1 cTG mice entered the anagen phase earlier than wild-type (WT) mice and showed the next hair coat by PND 63. Hair was fully regrown by PND 70 (Figure 1f). This difference in hair regrowth from the dermis to the skin surface was confirmed by hematoxylin and eosin (H&E) staining (Figure 1g). IF staining for the proliferation marker Ki67 revealed additional Ki67-positive cells in the HFs of AIMP1 cTG mice on PND 70 (Figure 1h).

Next, we generated systemically inducible AIMP1-transgenic mice (iTG) (Figure S2a). We confirmed that the mice were correctly generated by analyzing AIMP1 levels by using western blotting (Figure S2b). We clipped the dorsal skin of WT and AIMP1 iTG mice on PND 49 and observed hair cycle progression from telogen to anagen in the two types of mice under native conditions (Figure S2c). Four weeks later, AIMP1 iTG mice exhibited hair regrowth on over 80% of their back area, whereas most WT mice did not display hair regrowth (Figure S2d). AIMP1 iTG mice showed more anagen HFs than control mice (Figure S2e).

### Effect of AIMP1 peptide treatment on hair growth

To characterize the functional domain of AIMP1 involved in hair growth, we prepared four AIMP1 peptides based on a previous report (Figure S3a) ^19^. To determine the active region in hair regrowth, we used the four peptides of AIMP1 to treat DPCs and outer root sheath (ORS) follicular keratinocytes, which play important roles in hair biology. We further analyzed the accumulation of β-catenin in both cell types because β-catenin is a key molecule in hair growth and maintenance in the HF microenvironment ^24^. Full-length AIMP1 (FL), N192, and TN41, but not C120, elevated β-catenin levels in human DPCs (Figure S3b). By contrast, FL and TN41 did not increase β-catenin levels in the ORS cells.

We previously observed that AIMP1 inducible mice entered the anagen phase earlier, such that their hair regrowth rates differed from those of WT mice. To determine the telogen-to-anagen transition effect of TN41 on hair growth, we clipped the dorsal hair of C57BL/6 mice and applied vehicle and TN41 on the left and right sides of the clipped region, respectively (Figure S4a). After treatment with TN41 for 4 weeks, the TN41-treated regions exhibited more hair regrowth than the vehicle-treated regions (Figure 2a). Hair growth was verified using histological analysis. Additionally, we performed immunohistochemical staining to identify the expression of Ki67 in HFs. A higher staining intensity of Ki67 was observed in the secondary hair germ of TN41-treated mice 6 days after clipping (Figure 2b), and the Ki67 signal was observed in the area of the matrix cells (Figure S4b). To investigate the hair growth effects of TN41, we monitored how TN41 reached HFs using TN41 conjugated with a fluorescent dye and determined that TN41 flowed through the hair follicle pores (Figure S4c).

**Figure 2.**
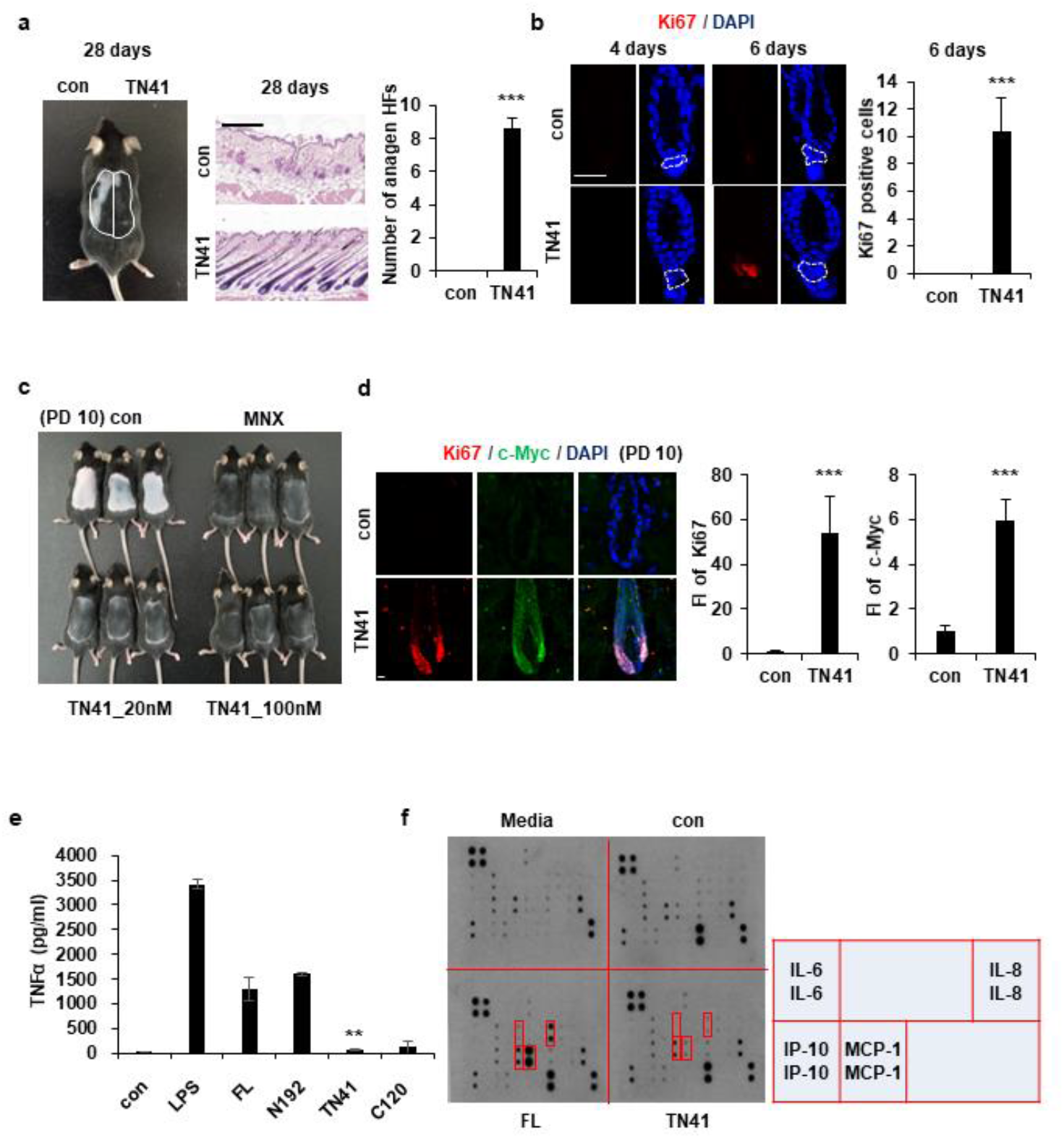
Effects of TN41 on hair growth. (**a**) Representative and H&E image at 28 days after clipping (n = 3). HFs area were measured. (**b**) IF images of Ki67 and counts of Ki67-positive cells. (**c**) Mice were depilated on PND 49 and treated with the vehicle, 3% MNX and TN41. Pictures were taken on PD10 (n = 3). (**d**) IF images for c-Myc and Ki67 and relative FI. (**e**) Effects of AIMP1 fragments on TNF-α secretion from RAW 264.7 cells. (**f**) Effects of FL and TN41 on cytokines secretion from DPC. Media: culture medium, con: cultured medium containing vehicle. The cytokines secreted by FL but not by TN41 are in red boxes. The bars: 200 μm (**a**) and 20 μm (**b** and **d**). Error bars: mean ± SD. ***: P < 0.001

To determine the effect of TN41 on hair growth rates, we depilated mice to synchronize the hair cycle to the beginning of the anagen phase. We treated depilated mice with TN41 and minoxidil (MNX). We used MNX, a hair growth-promoting compound, as a positive control for telogen-to-anagen transition. Hair was observed on the dorsal skin of mice in both the TN41-treated and MNX-treated groups, whereas hair growth was not apparent on the skin of vehicle-treated mice at 10 days post-depilation (PD10) (Figure 2c). Hair growth was verified using histological analysis (Figure S4d). We examined Ki67 and c-Myc levels in HFs using IF staining to confirm this result. Higher fluorescence staining of Ki67 and c-Myc was observed in TN41-treated HFs because TN41-treated HFs grew faster than vehicle-treated HFs (Figure 2d).

Interestingly, FL (but not TN41) caused signs of inflammation, such as skin swelling because of its known inflammatory activity (Figure S4e). To confirm this effect, we treated RAW 264.7 mouse macrophages with the four AIMP1 peptides and monitored the secretion of TNF-α, which is the signature cytokine of inflammation. FL and N192, but not TN41 or C120, induced the secretion of TNF-α, similar to lipopolysaccharide (Figure 2e). Next, we verified the presence of inflammatory cytokines in the DPCs after treatment with FL and TN41. FL, but not TN41, induced the secretion of inflammatory cytokines, including interleukin 6, interleukin 8, monocyte chemoattractant protein-1, and interferon gamma-induced protein 10 (Figure 2f). Therefore, to investigate the effects of AIMP1 on hair growth, which is uncoupled from its inflammatory function, we used TN41 in subsequent experiments.

### Mechanism of AIMP1 peptide in hair growth

A previous report showed that AIMP1 acts as a signaling molecule or an intracellular functional molecule ^25, 26^. Because the areas where TN41 was applied showed hair regrowth, we first confirmed whether TN41 functioned as a signaling molecule. AIMP1 was enriched in the bulge, with an expression pattern correlated with HFSCs (Figure S1c). Therefore, we evaluated the secretion of AIMP1 from HFSCs and the regulation of AIMP1 secretion by treating HFSCs with effector molecules. We observed cleaved AIMP1, approximately 20 kDa in size, in the Wnt3a-treated supernatant (Figure S5a). The cleaved form of AIMP1 was detected using western blot (WB) analysis using an antibody that binds to the N-terminal region of AIMP1 but not to its C-terminal region. To determine whether the detected peptide originated from AIMP1, we knocked down AIMP1 using an AIMP1-specific siRNA. The amount of secreted AIMP1 fragment decreased following treatment with siRNA against AIMP1 (Figure S5b).

A database-based peptide library was recently used to identify AIMP1 protease ^27^. AIMP1 is predicted to be a target candidate for matrix metalloproteinase (MMP). We conducted an *in vitro* cleavage assay to confirm whether AIMP1 was a target of MMPs. Among the MMPs, MMP1 cleaved AIMP1 within 2 h (Figure S5c). AIMP1 was completely cleaved after 4 h of incubation with MMP1. The cleaved form of AIMP1 was not detected when MMP1 was blocked with an inhibitor or antibody (Figure S5c). We performed a secretion assay with inhibitors of MMP1 to determine whether MMP1 influences the secretion of AIMP1 N-terminal peptides from cells. The secreted fragment of AIMP1 disappeared when the two types of inhibitors were co-treated with Wnt3a (Figure S5d). MMP1 is an extracellular protease. However, the intracellular activity of MMP1 has recently been reported. In particular, skin photo-aging is associated with the acceleration of collagen degradation by the intracellular activity of MMP1 ^28, 29, 30^. The cleaved peptide was identified using mass spectrometry and originated from the N-terminal sequence of AIMP1 (Figure S5e). These data support our hypothesis that TN41 is biologically valid.

Together with the β-catenin accumulation in DPCs via TN41 (Figure S3b), to identify the target cells of truncated AIMP1 *in vivo*, we used TN41 to treat the mouse dorsal skin and examined the expression level of lymphoid enhancer-binding factor 1 (LEF1) in DPCs. TN41-treated mice exhibited high LEF1 expression (Figure 3a). Ki67 and c-Myc were also highly expressed in DPCs following treatment with TN41 (Figure 3b). Additionally, β-catenin levels in DPCs were increased by TN41 in a concentration- and time-dependent manner (Figure 3c). Levels of *AXIN2* and *TCF7*, regulated by β-catenin, were also increased in DPCs via TN41 (Figure S6a). The expression of alkaline phosphatase, an established dermal papilla marker related to hair inductivity, was also increased by treating cells with TN41 in a dose- and time-dependent manner (Figure 3c).

**Figure 3.**
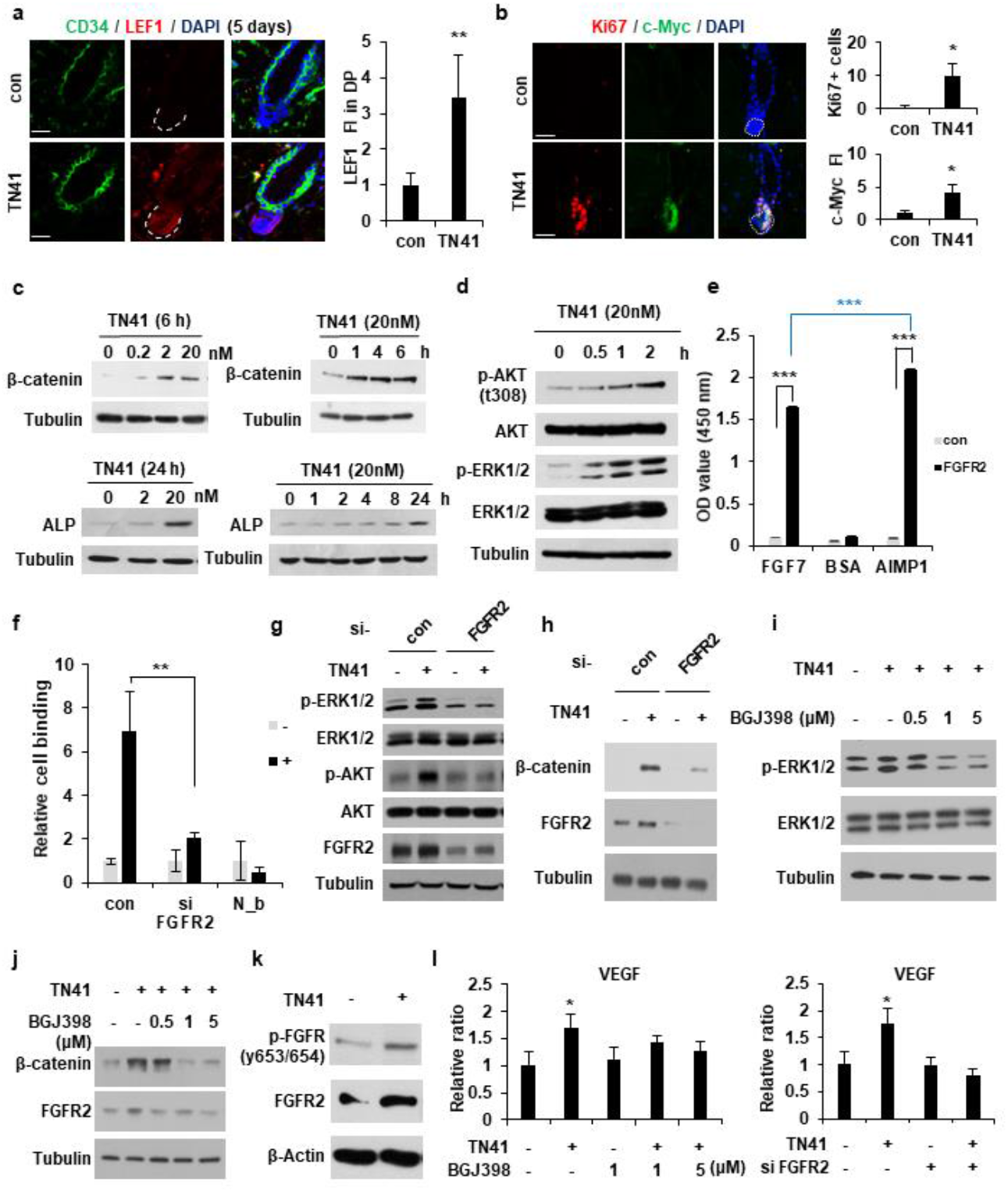
FGFR2-mediated activation of the MAPK pathway in DPCs by TN41. (**a**) IF images of CD34 and LEF1 from TN41-treated and vehicle-treated m. Relative FI in the DPC. (**b**) IF images of Ki67 and c-Myc. Ki67-positive cell counts and relative FI of c-Myc in the DPC. (**c and d**) DPCs were treated with TN41 and WB were performed. (**e**) Binding between AIMP1 and FGFR2. Gray bar: Fc (Fragment crystallizable), black bar: FGFR2-Fc. (**f**) FACS analysis to evaluate AIMP1 binding to DPCs. con: si-control, N_b: non-biotinylated TN41, -: untreated, +: TN41-treated. (**g - j**) TN41 was added to anti-FGFR2 siRNA-transfected and BGJ398-treated DPC. (**k**) TN41-induced phosphorylation of FGFR. (**l**) VEGF expression of TN41-treated DPCs with or without BGJ398 and anti-FGFR2 siRNA. The bars: 20 μm (**a** and **b**). Error bars: mean ±SD. ***: P < 0.001, **: P < 0.01, *: P < 0.05.

The N-terminal region of AIMP1 induces β-catenin expression in human bone marrow-derived MSCs via the FGFR2-mediated activation of Akt and ERK ^20^. We examined whether TN41 functions via a mechanism similar to that in MSCs of DPCs. Phosphorylation of Akt at residue 308 increased in a time-dependent manner following treatment with TN41 (Figure 3d). ERK phosphorylation revealed a pattern similar to that of Akt. We identified the functional receptor for TN41 in DPCs using chemicals that inhibit different target molecules posited to be involved in TN41-mediated DPC activation, including BGJ398 (FGFR1 and 2 inhibitors), erlotinib (epidermal growth factor receptor [EGFR] inhibitor), LDN-193189 (activin receptor-like kinase [ALK]2 and 3 inhibitors), crenolanib (platelet-derived growth factor receptor [PDGFR]a and b inhibitor), and repsox (ALK5 inhibitor). Among these, only BGJ398 blocked TN41 activity, which resulted in reduced β-catenin accumulation (Figure S6b). This suggests that FGFR2 is a potential receptor.

To test the interaction between TN41 and FGFR2, we tested AIMP1 and FGFR2 using ELISA and confirmed the interaction between the two molecules (Figure S6c). Moreover, AIMP1 revealed higher binding to FGFR2 than FGF7 (Figure 3e). The interaction between FGF7 and FGFR2 was inhibited by AIMP1, but not by bovine serum albumin (BSA; Figure S6d). The binding of TN41 to DPCs was decreased when FGFR2 was suppressed by its specific siRNA (Figure 3f). Although TN41 increased the phosphorylation levels of Akt and ERK, this effect was diminished when FGFR2 expression was suppressed by its specific siRNA (Figure 3g). β-catenin accumulation was also decreased by FGFR2 knockdown (Figure 3h). Phosphorylation of ERK was enhanced by TN41 but decreased after BGJ398 treatment (Figure 3i). The TN41-induced accumulation of β-catenin was ablated by treatment with BGJ398 (Figure 3j). Interestingly, TN41 enhanced FGFR2 levels (Figure 3g and h), which resulted in increased phosphorylation of FGFR (Figure 3k). Next, we evaluated the gene expression levels of secretory molecules in the DPCs. The expression levels of four positive genes involved in hair growth (*KGF*, *HGF*, *IGF*, and *VEGF*) were increased by TN41 (Figure S6e). However, the expression levels of four other inhibitory genes for hair growth (*TGFb1*, *TGFb2*, *DKK1*, and *IL6*) were not induced by TN41. The expression of *TGFb1* and *DKK1* was reduced. In accordance with these findings, TN41-mediated *VEGF* induction was reduced when FGFR2 expression was suppressed by BGJ398 and anti-FGFR2 siRNA (Figure 3l).

### Effect of TN41 on human hair shaft elongation in cultured HFs

We conducted a hair shaft elongation assay to determine whether AIMP1 had similar activity in human HFs. Because it was not possible to document the promotion of telogen-to-anagen activation in cultured human HFs, we decided to show that TN41 could also work on human HFs via the activation of DPCs. Human HFs were cultured in the presence or absence of TN41. After 6 days, we measured the length of the HFs. TN41-treated human HFs showed approximately 30% greater elongation than untreated controls (Figure 4a). Additionally, the number of Ki67-positive matrix keratinocytes around the DPCs was increased in TN41-treated HFs compared to untreated HFs (Figure 4b).

**Figure 4.**
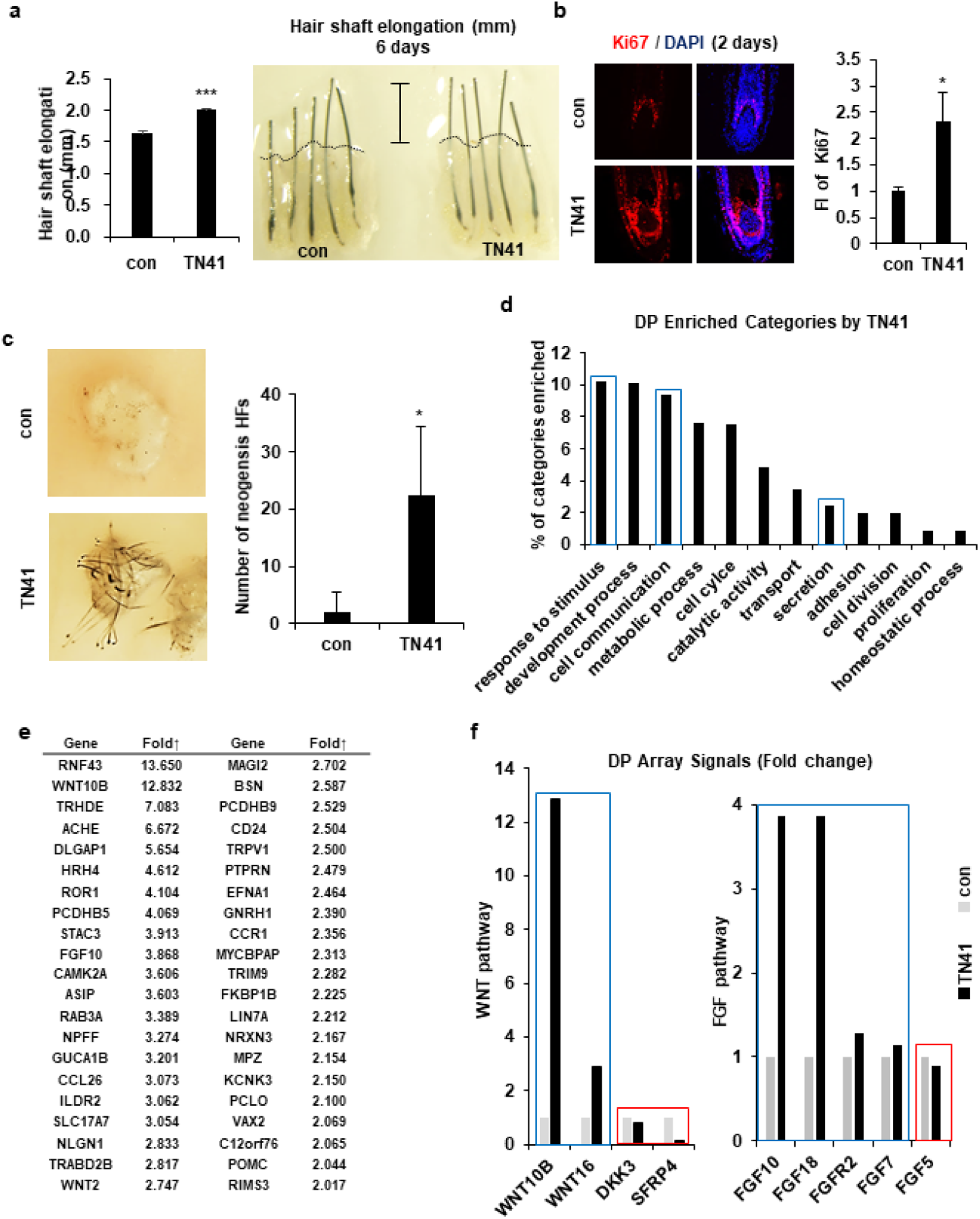
Effects of TN41 on human DPCs. (**a**) Human HFs were cultured for 6 days with TN41 or the vehicle and hair length was measured (3 donors). (**b**) IF image of Ki67 and relative FI. (**c**) 3D-cultured DPCs were injected into the hypodermis of nude mice and images were obtained after 2 weeks (injection site = 3). (**d**) Gene categories enriched in TN41-treated DPCs. Extracellular signaling molecules were among the most enriched (blue box). (**e**) List of DPC genes in extracellular signaling category that were upregulated. (**f**) The microarray signal values of Wnt and FGF pathway members displayed temporal differences. Blue box: positive genes, red box: negative genes. Error bars: mean ± SD. ***: P < 0.001, *: P < 0.05.

Next, we performed a patch hair reconstitution assay to examine whether TN41 could improve the hair-inductive activity of DPCs. 3D cultured human DPCs (100 spheres; 10^6^ cells) were treated with TN41 for 4 h and then co-transplanted with mouse epidermal cells (10^6^ cells) into the dorsal skin of nude mice. Hair induction increased after transplantation of TN41-treated human DPC spheres compared to that of untreated spheres (Figure 4c).

### TN41-mediated gene expression regulation in human DPCs

We compared the gene expression profiles of untreated and TN41-treated DPCs using microarray analysis, which revealed numerous differentially expressed genes between these groups. We derived the molecular signatures of genes with greater than two-fold upregulation or downregulation in TN41-treated DPCs relative to their untreated controls and performed bioinformatics analysis to classify these genes. The transcription of DPCs dynamically changed following treatment with TN41 (Figure 4d). Notably, the transcription of genes encoding putative hair cycle stimulating factors within the “extracellular signal” category was enhanced, and further analyses revealed elevated expression of 42 dermal papilla genes in the signaling category (Figure 4e). Interestingly, Wnt10b expression was highly upregulated by TN41. Wnt10b promotes the growth and regeneration of HFs via the canonical Wnt signaling pathway ^31^. Next, we focused on the Wnt signaling pathway because Wnt2 and Wnt10b were upregulated by TN41 (Figure 4f). Additionally, Wnt16 activates human keratinocyte proliferation and differentiation via the non-canonical Wnt transduction pathway ^32^. DKK3 and SFRP4 are potent inhibitors of the Wnt signaling pathway ^33, 34^. *FGF7* and its relative gene *FGF10* were identified as DPC signature genes elevated by TN41 (Figure 4e and f). FGF7 and FGF10 expression in DPCs coincided with an increase in MAPK pathway activity (Figure 3d) ^35^. Together with the results of previous studies, this suggests a paracrine role for FGF7 and/or FGF10 signaling in epidermal wound repair and hair growth ^36, 37^. FGFR2 showed the same trend, consistent with previous data (Figure 3k). FGF18 expression was higher in TN41-treated DPCs than in untreated DPCs. This trend for FGF18 was consistent with its anagen-inducing effects on DPCs ^38^. By contrast, FGF5 showed the opposite trend, consistent with its growth-inhibitory effects on DPCs ^39^.

## DISCUSSION

Recent studies have revealed the complexity of cellular and molecular regulators within the skin stem cell niche during development, homeostasis, injury, and aging. Although many putative niche factors have been identified, crosstalk between HFSCs and DPCs remains unclear. In this study, we identified the function of AIMP1 in the HF environment. Wnt-activated HFSCs secrete truncated N-terminal AIMP1, which acts as a ligand for FGFR2 in DPCs, thereby activating the MAPK pathway; this, in turn, leads to the accumulation of β-catenin in DPCs. These processes may activate the secretion of anagen-inducing molecules from DPCs. Our results are consistent with data showing that AIMP1 enhances the proliferation of bone marrow-derived MSCs via the FGFR2-mediated accumulation of β-catenin ^20^. Homeostatic signals governing the connections between HFSCs and the surrounding cells remain unclear ^40, 41^; therefore, additional studies on AIMP1 are needed to provide further insights.

Communication between HFSCs and DPCs leads to physical structural development, maintenance, and hair follicle cycling. Soluble factors derived from these two cells contribute to microenvironmental changes during the asynchronous cycle. Ensuring the balance between apoptosis-associated factors, inducing the anagen-to-catagen transition via apoptosis and growth-associated factors, and maintaining the anagen phase through growth promotion are important tasks. Alopecia-inducing factors, including hormones, inflammatory cytokines, and senescence molecules, disrupt communication between HFSCs and DPCs by inducing apoptosis or senescence of HFSCs and DPCs. Growth-associated factors are potential drug targets. Although growth factors are viable drug candidates, they do not have sufficient stability. However, TN41 demonstrated a more stable structure ^42^ than the AIMP1 protein and showed hair growth-promoting activity through the activation of DPCs. Moreover, TN41 was suitable for peptide synthesis. Based on these features, we expect TN41 or TN41-derived smaller peptides to be potential drug candidates for addressing hair loss.

Overall, our results demonstrated that AIMP1 participates in the maintenance of HFs, especially by promoting hair cycle transition from the telogen phase to the anagen phase. Because TN41 enhances hair growth rate and telogen-to-anagen transition in a mouse model and hair shaft elongation in cultured human HFs, this peptide may mitigate hair loss. Moreover, the enhanced hair-inducing activity of cultured human DPC spheroids after TN41 treatment in a hair reconstitution assay suggests that AIMP1 may also be useful as an adjuvant in cell-based therapy to overcome hair loss.

## MATERIALS AND METHODS

### Generation of inducible hAIMP1 mice

For generation of the inducible hAIMP1 knockin mouse, the tetO-hAIMP1 construct was cloned to contain human AIMP1 cDNA under the control of a minimal promoter from hCMV fused to the tetO sequence. This construct was subcloned into a ROSA targeting vector (Soriano P’s lab, NY, USA). The targeting vector was electroporated into mouse embryonic stem (ES) cells (E14TG2a). Correctly targeted clones were injected into C57BL/6 blastocysts for chimera generation. TRE-hAIMP1 mice were crossed to CAG-rtTA3 (Jackson Laboratories, Bar Harbor, ME), KRT14-rtTA (Jackson Laboratories), or KRT14-cre (Jackson Laboratories) with Rosa26-CAGs-LSL-rtTA3 (Jackson Laboratories) to generate systemic or skin-specific hAIMP1-inducible mice.

### Immunofluorescence staining

Paraffin sections were used for IF analysis. Slides were incubated for 15 min at 60℃, and immersed in xylene twice for 5 min for deparaffinization. Slides were soaked in 100% EtOH twice for 2 min, 95% EtOH twice for 1 min, and 80% EtOH, 70% EtOH, and deionized H2O for 1 min each for rehydration. Antigen retrieval was performed by boiling the slides for 20 min in antigen unmasking solution (Vector Laboratories, Burlingame, CA, USA). Slides were cooled in buffer for 15 min, rinsed gently with running water for 5 min, and washed with PBS for 5 min. Nonspecific binding was blocked by incubation with CAS block (Life Technologies, CA, USA) for 30 min. Tissue sections were incubated with primary antibodies at 4° C overnight and then washed three times with PBS containing 0.2% Triton X-100 (PBSTx; Sigma-Aldrich, St. Louis, MO). The following antibodies were used: CD34 (Abcam, Cambridge, UK), Ki67 (eBioscience, San Diego, CA), beta-catenin (Abcam), LEF1 (Thermo Fisher Scientific, Waltham, MA), c-MYC (Abcam), KRT15 (Abcam), SOX9 (eBioscience), GFP (Invitrogen, Carlsbad, CA), AIMP1 (Thermo Fisher Scientific), and EPRS (Thermo Fisher Scientific). Sections were incubated with secondary antibodies conjugated with Alexa Fluor 488 or 594 (Invitrogen). After washing three times with PBSTx, 4′, 6-diamidine-2′-phenylindole dihydrochloride (Thermo Fisher Scientific) was added for nuclear counterstaining. Coverslips were mounted onto glass slides with fluorescent mounting medium (Biomeda Corp., Burlingame, CA). All images were obtained by A1 confocal microcopy (Nikon, Japan).

### Animals

C57BL/6 mice were purchased from Orient bio (Sungnam, Korea). Offspring were genotyped by PCR-based assays of mouse tail DNA. Animal care was conducted in accordance with the guidance of Seoul National University. All animal experiments were performed following the Guidelines for the Care and Use of Laboratory Animals.

### Western blotting

Cells were harvested with lysis buffer (25 mM Tris-HCl, pH 7.5, 100 mM NaCl, 5% glycerol, 0.5% Triton X-100, and 1 mM EDTA) containing a 1x protease inhibitor tablet (Roche, Basel, Switzerland) and a 1x phosphatase inhibitor tablet (Roche). The protein concentration was measured by Bradford assay. Whole-cell lysates were subjected to gel electrophoresis. Using the semi-dry transfer method, proteins were transferred to polyvinylidene fluoride membranes. The membranes were immersed in 5% skim milk (BD Biosciences, Franklin Lakes, NJ, USA) for 30 min and then incubated with each primary antibody diluted in 1% skim milk. Primary antibodies used in this study were alkaline phosphatase (ALP; Santa Cruz Biotechnology, Dallas, TX, USA), beta-catenin (Cell Signaling Technology, Danvers, MA, USA), N-AIMP1 (Atlas Antibodies, Bromma, Sweden), C-AIMP1 (Novus Bio, Littleton, CO, USA), Akt (Cell Signaling Technology), phospho-Akt (Cell Signaling Technology), ERK (Cell Signaling Technology), phospho-ERK (Cell Signaling Technology), FGFR2 (Abcam, Cambridge, UK), phosphor-FGFR (cell signaling), MMP1 (Thermo Fisher Scientific), beta actin (Santa Cruz Biotechnology) and tubulin (Novus Bio). After washing three times for 5 min each with 1x Tris-buffered saline containing Tween 20 (0.1% TBST), the membranes were incubated for 1 h with horseradish peroxidase (HRP)-conjugated secondary antibodies diluted in 1% skim milk and washed three times for 5 min each with 0.1% TBST. After incubation with western HRP substrate detection solution (Abcam), immunoblot images were acquired.

### In vivo ‘patch’ hair reconstitution assay

‘Patch’ hair reconstitution assays were conducted in this study. To isolate mouse dermal cells, the dorsal skin of C57BL/6 neonates was collected and incubated overnight with 1 mg/mL collagenase/dispase (Roche). The dermis and epidermis were separated by incubating the skin with 0.25% trypsin/10 mM EDTA in phosphate-buffered saline (PBS) at 37℃ for 15 min. Epidermal and dermal cells were filtered through cell strainers of 70 and 100 µm (BD Biosciences) respectively, and then centrifuged at 1500 rpm for 5 min. Three-dimensional (3D) DPCs were combined with freshly isolated neonatal mouse epidermal cells (1 × 106 cells) and then subcutaneously co-implanted into the skin on the backs of 7-week-old female nude mice. After 2 weeks, back skin was collected from the mice, and reconstituted HFs were quantified.

### In vitro cleavage assay

AIMP1 and MMP1,2 and 9 (Novus Bio) were incubated in cleavage assay buffer (20 mM HEPES pH 7.4, 140 mM NaCl, 2 mM CaCl2) for 4 h at 37°C. ARP100 (Santa Cruz Biotechnology), doxycycline hyclate (Sigma-Aldrich, St. Louis, MO, USA) and anti-MMP1 antibody (R&D Systems, Minneapolis, MN, USA) were used for each assay. Proteins were separated by sodium dodecyl sulfate polyacrylamide gel electrophoresis (SDS-PAGE) and subjected to immunoblotting.

### Hair depilation and peptide administration

For hair cycle induction, hair on the dorsal skin of 7-week-old mice was depilated to induce a new hair cycle after confirming that the skin had a light pink color, which indicates that the HFs are synchronized at the telogen phase. For peptide treatment, 150 µ L of peptides in carbomer (20 and 100 nM) were topically applied to the depilated area once per a day. We used thioglycolate hair removal cream (Ildong Pharmaceutical, Seoul, Korea) for chemical depilation.

### Cell binding assay by fluorescence-activated cell sorting (FACS)

TN41 was biotinylated with an EZ-LinkTM Sulfo-NHS-LC-Biotinylation kit (Thermo Fisher Scientific) following the manufacturer’s instructions. DPCs (1.8 × 105) were cultured in six-well plates containing Dulbecco’s modified Eagle’s medium (DMEM; Gibco BRL, Grand Island, NY, USA) with 10% fetal bovine serum (FBS), 1 ng/mL of FGF2 (Abcam), and 1% antibiotics for 24 h and then treated with biotinylated 1 µM TN41 for 1 h. The cells were detached by Accumax (Merck Millipore, Billerica, MA, USA) and washed twice with PBS containing 1% bovine serum albumin and 0.05% NaN3 (FACS buffer). Cells were fixed by 4% paraformaldehyde for 10 min and then washed once with FACS buffer. The cells were incubated with phycoerythrin-conjugated streptavidin antibody (R&D Systems) diluted in FACS buffer for 30 min and then washed twice with FACS buffer. Cell were resuspended in FACS buffer and measured with a BD accuriTM C6 plus flow cytometer (BD Biosciences).

### Binding by enzyme-linked immunosorbent assay (ELISA)

Microplates were coated with 0.5 μM of human IgG1 Fc, FGFR2-Fc (Curebio, Suwon, Korea) in PBS for 1 h at 37 °C. Microplates were washed with PBS containing Tween 20 (0.05 % PBS-T) three times. Plates were blocked with 3 % BSA in 0.05% PBS-T for 1 h at 25 °C. 1 μM of biotinylated AIMP1 (Curebio) was added into plates and incubated for 1 h at 25 °C. Streptavidin HRP (Genescript, Piscataway, NJ, USA) was added and incubated for 1 h at 25 °C. TMB solution (Thermo Fisher scientific) was applied for 3 min and the reaction was stopped with 2N H2SO4. Signals were obtained with microplate reader (Biotek, Winooski, Vermont, USA) at OD 450 nm.

### Binding comparison by ELISA

A Maxisorp plate (Nunc, Roskilde, Denmark) was coated with recombinant proteins, AIMP1 (CureBio, Seoul, Korea), FGF7 (Miltenyi Biotec, Bergisch Gladbach, Germany), and BSA (Thermo Fisher Scientific) for 15 h at 4°C. For blocking, the coated plate was incubated with 5% bovine serum albumin in PBS. After blocking, FGFR2-flag recombinant protein (Origene, Rockville, MD, USA) was added to the plate for adherence to the coated binding candidates. After washing with PBS-T (0.05% Tween-20), the plate was incubated with anti-flag M2 antibody (Sigma-Aldrich). The plates were further incubated with anti-mouse IgG HRP (Merck Millipore). To develop the signal, 1-Step Ultra TMB ELISA substrate (Thermo Fisher Scientific) was added. The absorbance at 450 nm was measured with a microplate reader (TECAN, Männedorf, Switzerland).

### Competition assay by ELISA

Microplates were coated with 0.5 μM of FGF7 (R&D Systems) in PBS for 1 h at 37 °C. Microplates were washed with 0.05 % PBS-T three times. Plates were blocked with 3 % BSA in 0.05 % PBS-T for 1 h at 25 °C. 1 μM of FGFR2-Fc (Curebio) was pre-incubated with 0 ∼ 10 μM of BSA (Thermo fisher scientific) or 1 ∼ 10 μM of AIMP1 (Curebio) for 1 h at 25 °C then samples were incubated with FGF7 for 1 h at 25 °C. Anti-Human HRP (Genescript) was added and incubated for 1 h at 25 °C. TMB solution (Thermo fisher scientific) was applied for 5 min and the reaction was stopped with 2N H2SO4. Signals were obtained with microplate reader (Biotek) at OD 450 nm.

### Isolation and culture of human HFs

The Medical Ethical Committee of Kyungpook National University (Daegu, Korea) approved all described experiments (IRB No. KNU-2021-0113). Occipital scalp biopsy specimens were obtained during hair transplantation in human male patients with androgenetic alopecia. The Medical Ethical Committee of Kyungpook National University Hospital (Daegu, Korea) approved all described experiments. HFs were isolated from non-balding scalps as previously described with minor modifications ^43, 44^. Briefly, the subcutaneous fat portion of the scalp skin, including the lower HFs, was dissected from the epidermis and dermis. HFs were then isolated under a binocular microscope using forceps. Isolated HFs were maintained in Williams E media (Sigma-Aldrich) supplemented with 2 mM L-glutamine, 100 U/mL streptomycin, and 10 ng/mL hydrocortisone. Follicles were maintained in a humidified atmosphere of 5% CO_2_ at 37 °C.

### Cell culture

The dermal papilla was isolated from the bulbs of dissected HFs, transferred onto plastic dishes coated with bovine type I collagen, and cultured in DMEM supplemented with 1% antibiotic-antimycotic solution and 20% heat-inactivated FBS at 37°C in a humidified atmosphere containing 5% CO_2_. These explants were incubated for several days, and the medium was changed every three days. Once sub-confluent, the cells were harvested with 0.25% trypsin/10 mM EDTA in PBS, split at a 1:3 ratio, and maintained in DMEM supplemented with 10% FBS and 1ng/mL of FGF2. Non-balding scalp specimens were obtained from patients undergoing hair transplantation surgery. The ORS cells isolated from HFs were cultured as described before ^45^. Briefly, the hair shaft and hair bulb regions of the HF were cut off to prevent contamination with other cells. Trimmed HFs were immersed in DMEM supplemented with 20% fetal bovine serum. On the third day of culture, the medium was changed to keratinocyte growth medium (Gibco BRL) containing penicillin, streptomycin, and fungizone. After subculture, cells were maintained in keratinocyte growth medium and cells from the second passage were used in this study. HFSCs were purchased from Cellprogen (Torrance, CA, USA). HFSCs were used before passage six.

### Reverse transcription PCR and real-time PCR

Total RNA was isolated from DPCs by using the RNeasy Mini Kit (Qiagen, Hilden, Germany), according to the manufacturer’s protocol. Complementary cDNA was synthesized by RevertAid First Strand cDNA Synthesis Kit (Thermo Fisher Scientific), according to the manufacturer’s protocol. The real-time PCR was performed using Step one Plus real time PCR Assay (Applied Biosystems, Foster City, CA). All reactions were carried out using Power SYBR Green premix (Applied Biosystems).

### TNF-α ELISA assay

RAW264.7 mouse macrophages (2 × 104) were cultured in 24-well plates containing DMEM with 10% FBS and 1% antibiotics for 12 h and then treated with the purified protein for 6 h. The media were harvested after centrifugation at 3,000 ×g for 5 min, and secreted TNF-α was measured using a TNF-α ELISA kit (BD Biosciences) following the manufacturer’s instructions.

### Mass spectrometry

GST-AIMP1 was in-gel digested with Trypsin GOLD (Promega, Madison, WI). Tryptic peptides were separated by reversed-phase chromatography using Eazynano LC II (Thermo Fisher Scientific) with EASY-Column (100 μm inner diameter, 2 cm length) and EASY-Column (75 μm inner diameter, 10 cm length, 3 μm particle size, both Thermo Fisher Scientific). The separated peptides were analyzed using LTQ-Orbitrap Velos mass spectrometer (Thermo Fisher Scientific) system in positive ion mode. Electrospray (ESI) voltage was +1.9 kV and capillary temperature was 275°C. Spectra were acquired in a full scan mass (range 300 – 2,000 m/z) and followed by data-dependent collision-induced dissociation (CID) MS/MS scans. The MS/MS spectra were searched against GST-AIMP1 sequence database using Proteome Discoverer (version 1.3, Thermo Fisher Scientific) with SEQUEST search engine. Precursor mass tolerance and fragment mass tolerance were set to 25 ppm and 0.8 Da. Methionine oxidation (+15.99 Da) was set as variable modifications and cysteine carbamidomethylation (+57.02 Da) was set as static modification.

### Microarray analysis

Total RNA was extracted with the RNeasy Micro Kit. The Illumina NextSeq500 array (San Diego, CA, USA) of EBiogen (Seoul, Korea) was used. To analyze the Gene Ontology (GO) terms with ≥2.0-fold increased or decreased gene clusters, the web-based Database for Annotation, Visualization and Integrated Discovery (DAVID) 6.7 was used. To extract reliable GO terms belonging to biological processes, Fisher’s exact test and multiple test correction (p < 0.05) were used.

### Secretion confirmation by western blotting

HFSCs were cultivated to 60% confluence in DMEM containing 10% FBS. Cells were washed twice, and transferred to serum-free DMEM for 2 h. Cells were treated with Wnt3a (R&D Systems), Noggin (Miltenyi Biotec), FGF7 (Sigma-Aldrich), sonic hedgehog (SHH) (Miltenyi Biotec), or transforming growth factor (TGF)-β2 (Thermo Fisher Scientific) followed by incubation for 24 h. The culture media were collected and centrifuged at 500 ×g for 10 min. Supernatants were centrifuged again at 10,000 ×g for 30 min to eliminate membrane organelles, and the proteins were precipitated with 12% trichloroacetic acid and incubated for 12 h at 4°C. Pellets were obtained by centrifugation at 18,000 ×g for 15 min and neutralized with 100 mM HEPES, pH 8.0. Protein in the pellets were separated by SDS-PAGE and subjected to immunoblotting.

### Preparation of paraffin and frozen sections

For paraffin sections, mouse skin or human HF was fixed in 10% formalin solution at 4°C overnight and then embedded in paraffin. Paraffin-embedded specimens were cut into 5 µm-thick sections. For frozen sections, mouse skin or human HF tissues were immersed in ice cold 4% paraformaldehyde in PBS (pH 7.4) for 10 min. The fixed samples were embedded in optimal cutting temperature compound (Sakura FineTechnical Co., Ltd), snap-frozen in liquid nitrogen, and stored at –80°C. Frozen samples were cut in 10 µm-thick sections.

### Hematoxylin and eosin staining

Paraffin sections were deparaffinized and then rehydrated. Frozen sections were removed from the optimal cutting temperature compound and stained with hematoxylin and eosin (H&E; Sakura FineTechnical Co., Ltd., Tokyo, Japan) for tissue histology analysis using an optical microscope.

### Cytokine array

DPCs were cultured in 12-well plates containing DMEM with 10% FBS, 1 ng/mL of FGF2, and 1% antibiotics for 24 h and then treated with the AIMP1 FL (20 nM) and TN41 (20 nM) for 12 h. Effect of AIMP1 FL and TN41 on the secretion of different cytokines was determined using cytokine array (R&D Systems) following the manufacturer’s instructions.

## DATA AVAILABILITY STATEMENT

RNA sequencing data can be acquired upon successful application at Dryad (https://doi.org/10.5061/dryad.v41ns1rzb). All remaining data are available in the manuscript or supplementary materials.

## ACKNOWLEDGMENTS

SK

This research was supported by the Yonsei University Research Fund of 2020-22-0358, 2020-22-0356, 2021-22-0061 and NRF-2021R1A3B1076605 from the National Research Foundation of Korea.

YKS

This research was supported by the Basic Science Research Program through the National Research Foundation of Korea (NRF-2021R1A2C2003875) and Commercializations Promotion Agency for R&D Outcomes (COMPA) grant funded by the Korea government (MSIT) (No. 2021N100).

## AUTHOR CONTRIBUTIONS

Conceptualization: YK; Methodology: YK; Formal analysis: YK, HL, DK, SSB, IY, SBK, SC, SJJ, YJ. JWO, JMP, MCP; Writing - original draft preparation; YK; Writing - review and editing: YK, RP, YKS, SK; Supervision: SK

## SUPPLEMENTARY FIGURES

**Supplementary Figure 1.**
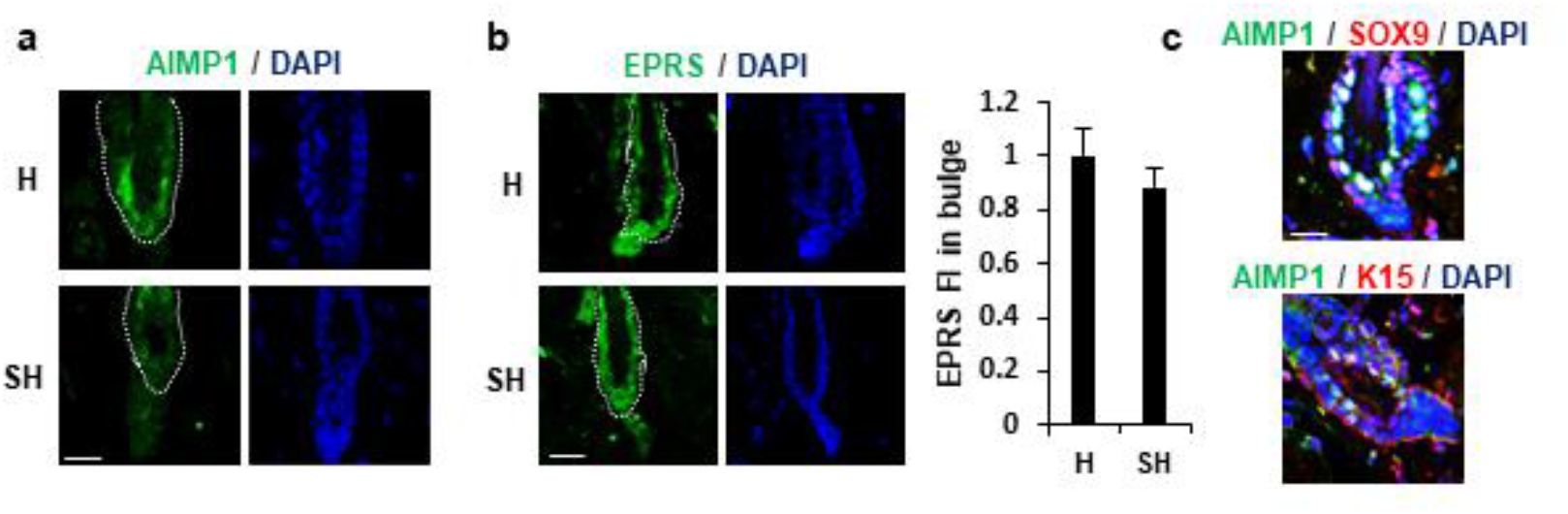
Positive correlation of AIMP1 with hair follicle maintenance. (**a**) IF images of AIMP1 from H and SH mice. Bulge was surrounded by dotted line. (**b**) IF images of EPRS from H and SH mice. Bulge was surrounded by dotted line. Relative FI of EPRS in the bulge was measured using Image J software (**c**) IF images of AIMP1, SOX9, and K15 in the bulge region from 2-month-old C57BL/6. The scale bars indicate 20 μm (**a**, **b** and **c**). Error bars indicate mean +/- SD. ***: P < 0.001.

**Supplementary Figure 2.**
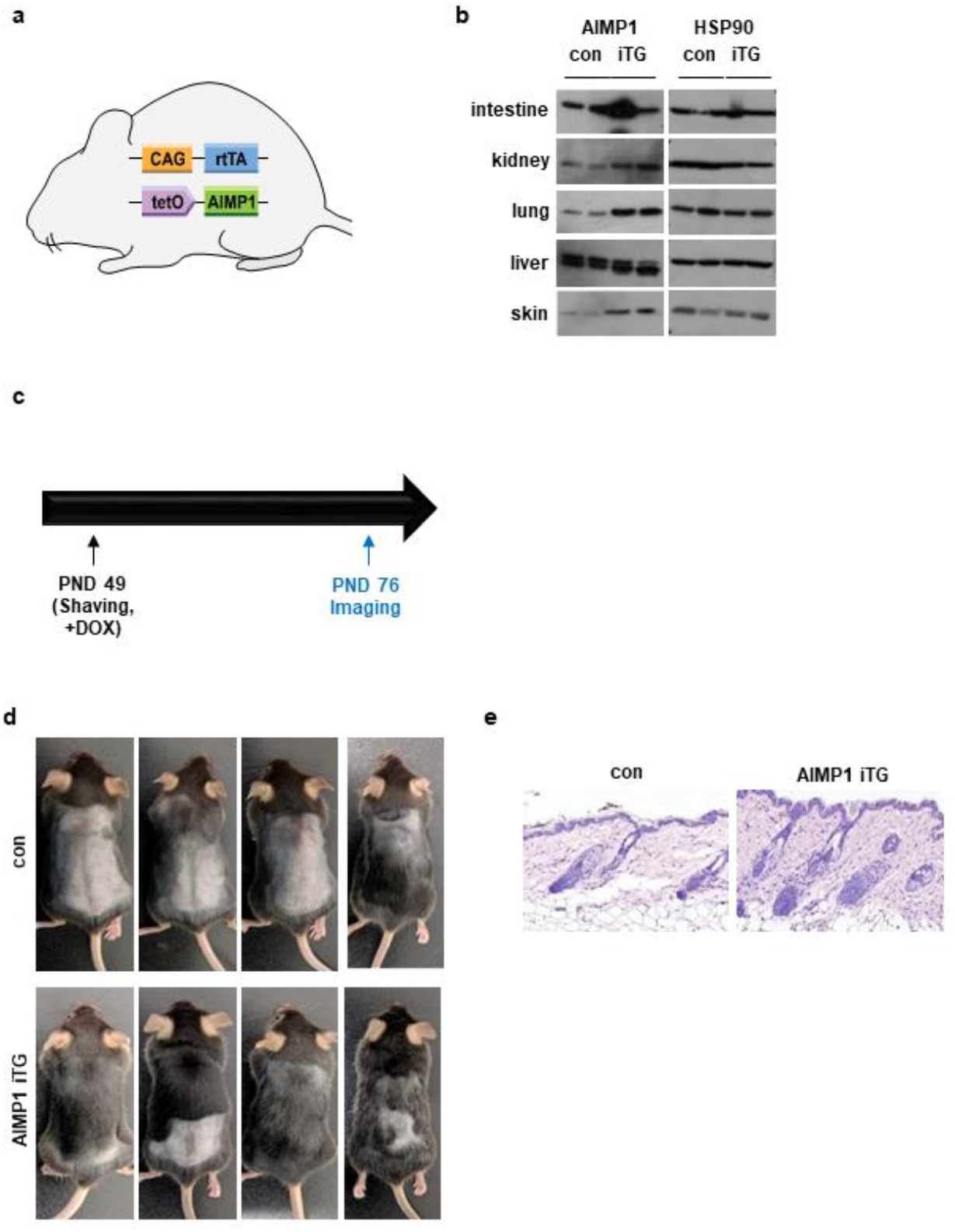
Identification of AIMP1 function with AIMP1-induced mice. (**a**) A schematic diagram of systemic-human AIMP1-inducible mouse model, CAG-rtTA; tetO-hAIMP1 (iTG). (**b**) Characterization of iTG mice by WB and immunohistology analysis. AIMP1 levels between the WT and iTG mice were compared by WB in the indicated organs. Induction of HA-tagged AIMP1 was determined by IF staining with an anti-HA antibody. (**c**) The experimental design for analysis of hair growth. WT and iTG mice were clipped at postnatal 49 days. 1.6 mg/mL of doxycycline contained water was supplied at the same day. Images were obtained at PND 49 and PND 76. (**d**) Mouse images of WT (con) and iTG. (**e**) H&E stained tissue images in WT and iTG mice.

**Supplementary Figure 3.**
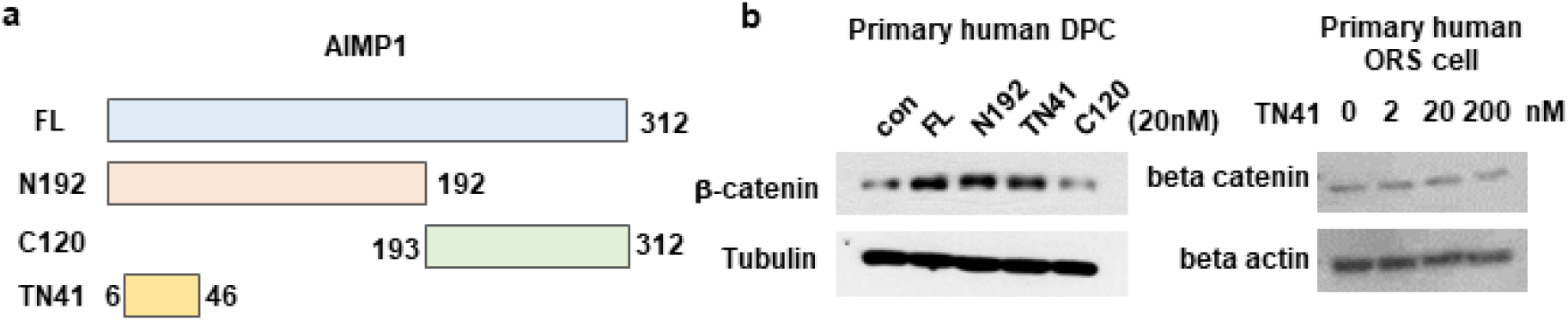
Determination of the peptide region of AIMP1 responsible for the hair growing activity. (**a**) Fragments of AIMP1. FL: Full length, N192: 1-192aa, C120: 193-312aa, TN41: 6-46aa. (**b**). Human primary DPCs were treated with FL, N192, TN41, or C120 (20 nM each). WB were performed with an anti-beta-catenin antibody. Human primary ORS cell were treated with various concentration of TN41.

**Supplementary Figure 4.**
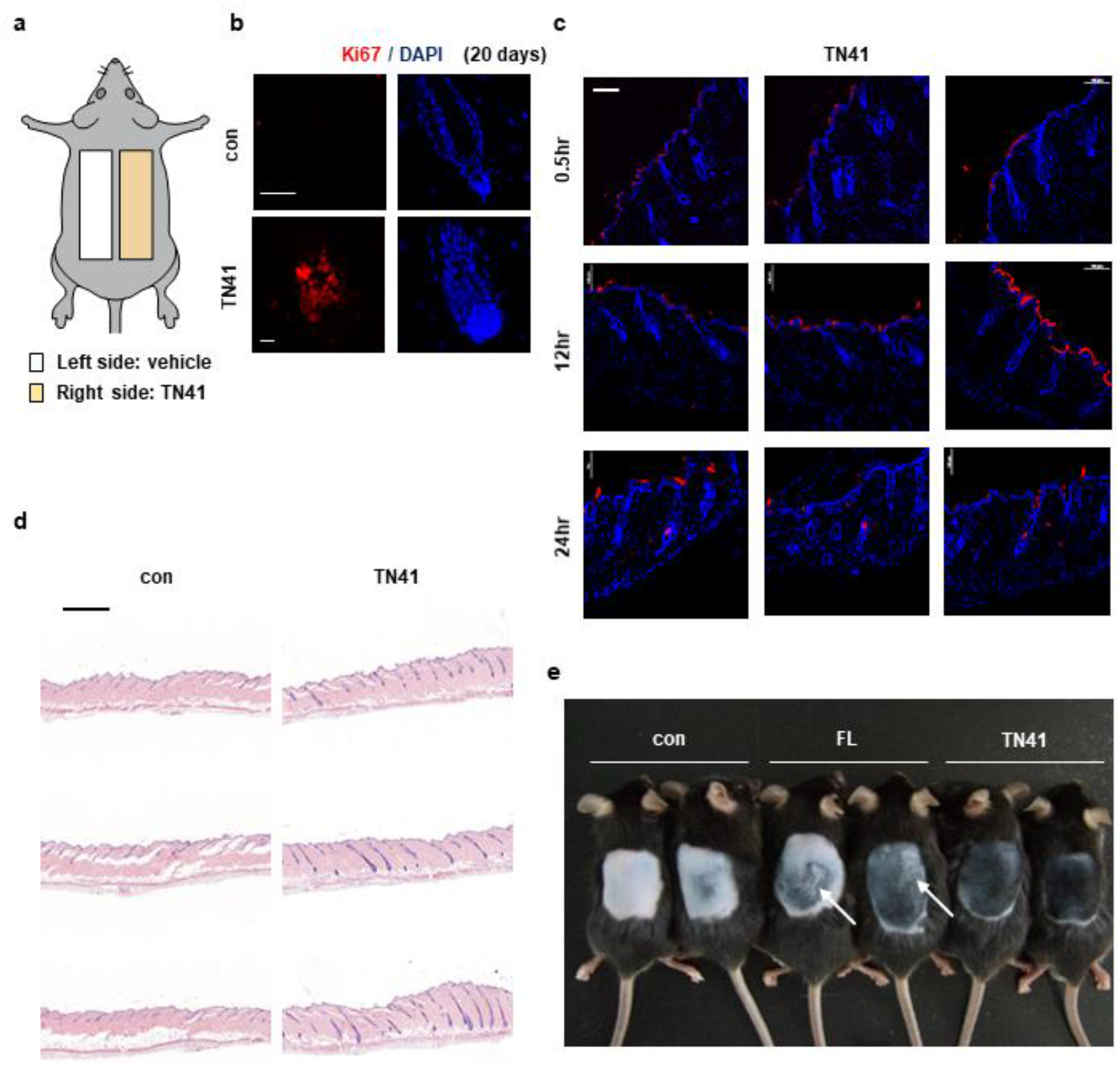
Effect of TN41 treatment on hair growth. (**a**) Mouse back skins were clipped at PND 49. The left and right halves of the back skin were treated with vehicle, TN41 (100 nM), respectively. (**b**) IF images with Ki67 from TN41-treated and non-treated region at 20 days after clipping. (**c**) Alexa 647-conjugated TN41 was applied to the back skin of mice. The skin samples were taken at the indicated time to make the frozen block. The fluorescence images were taken by A1 confocal microscopy. Red: Alexa 647 signal, Blue: DAPI. (**d)** H&E image of the treated and untreated dorsal skin. (**e**) Mouse back skins were depilated at PND 49, and images were taken PD10. White arrows indicate the skin regions showing inflammatory response. The scale bars indicate 20 μm (**b**), 100 μm (**c**), and 500 μm (**d**).

**Supplementary Figure 5.**
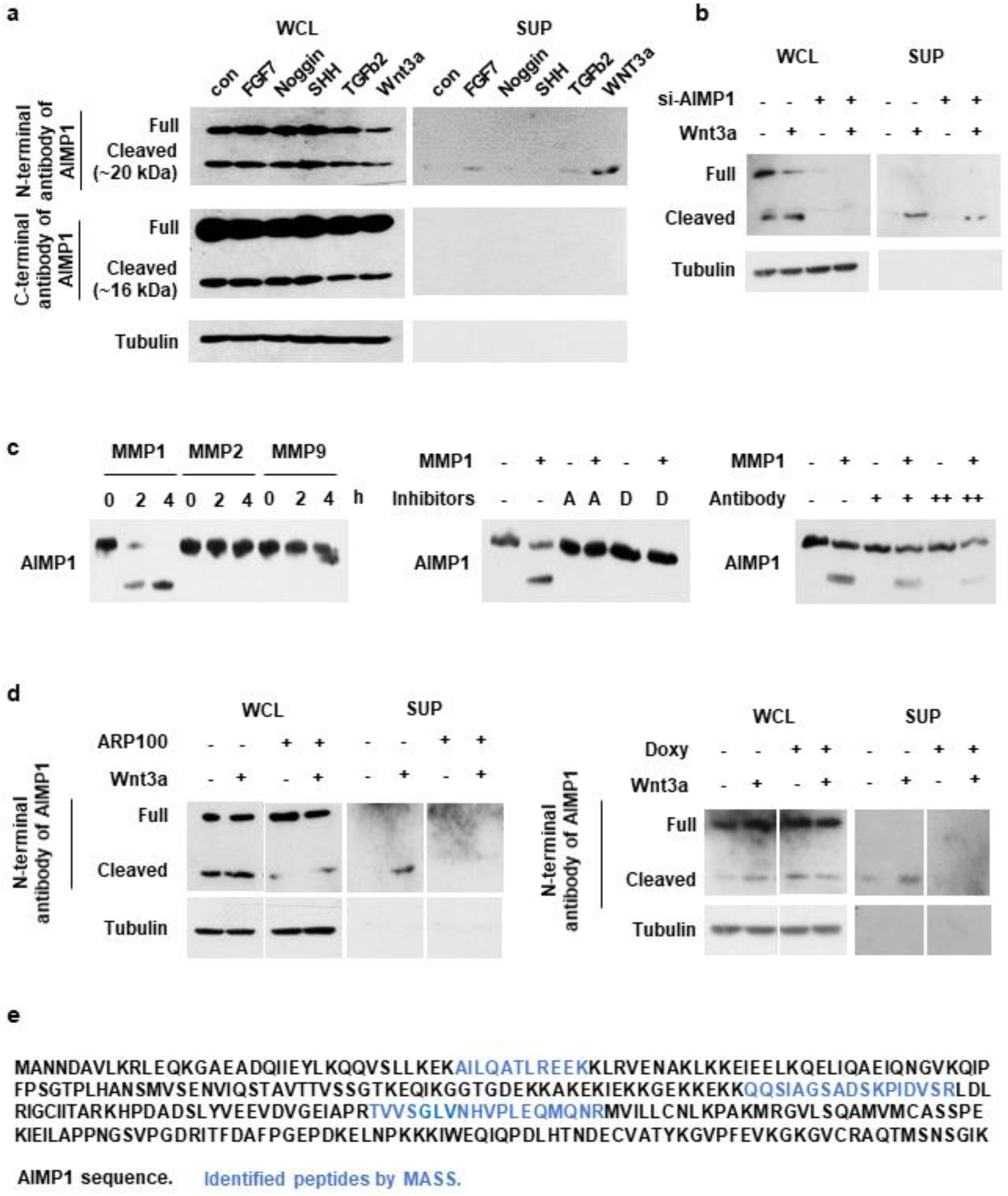
Secretion of AIMP1 N-terminal fragment from Wnt-activated HFSCs. (**a**) After serum starvation for 2 h, HFSCs were treated with indicated signaling molecules and incubated for 24 h. The proteins in the whole cell lysates (WCL) and supernatant (SUP) were subjected to western blot analysis with the antibodies specific to the N-terminal and C-terminal regions of AIMP1. Wnt3a: 200 ng/ml, FGF7: 100 ng/ml, Noggin: 200 ng/ml, TGFβ2: 10 ng/ml, SHH: 200 ng/ml. (**b**) AIMP1 siRNA (10 pmol) was transfected into HFSCs for 48 h before secretion assay. After serum starvation for 2 h, HFSCs were treated with 200 ng/ml of Wnt3a for 24 h and western blot was performed with the N-terminal specific AIMP1 antibody. (**c**) AIMP1 was incubated with each of MMP1, MMP2 and MMP9 at 37 °C for 4 h and western blot analysis with the N-terminal specific AIMP1 antibody. AIMP1 was incubated with MMP1 in the absence and presence of ARP100 (A, MMP1 and MMP2 inhibitor) and doxycycline hyclate (D, MMP1, MMP9 and MMP12 inhibitor). AIMP1 was incubated with different amounts of an anti-MMP1 antibody. AIMP1 (400 ng), MMP1 (72.9 ng), MMP2 (100 ng), MMP9 (106 ng), ARP100 (1.5 μg), doxycycline hyclate (1 μg), and anti-MMP1 antibody (+: 1 μg and ++: 1.5 μg). (**d**) Secretion of AIMP1 peptide fragment was confirmed by western blot. The band of secreted AIMP1peptide fragment was disappeared by MMP1 inhibitors. (**e**) The AIMP1 peptide fragments resulted from the MMP1 cleavage assay were identified by mass spectrometry analysis.

**Supplementary Figure 6.**
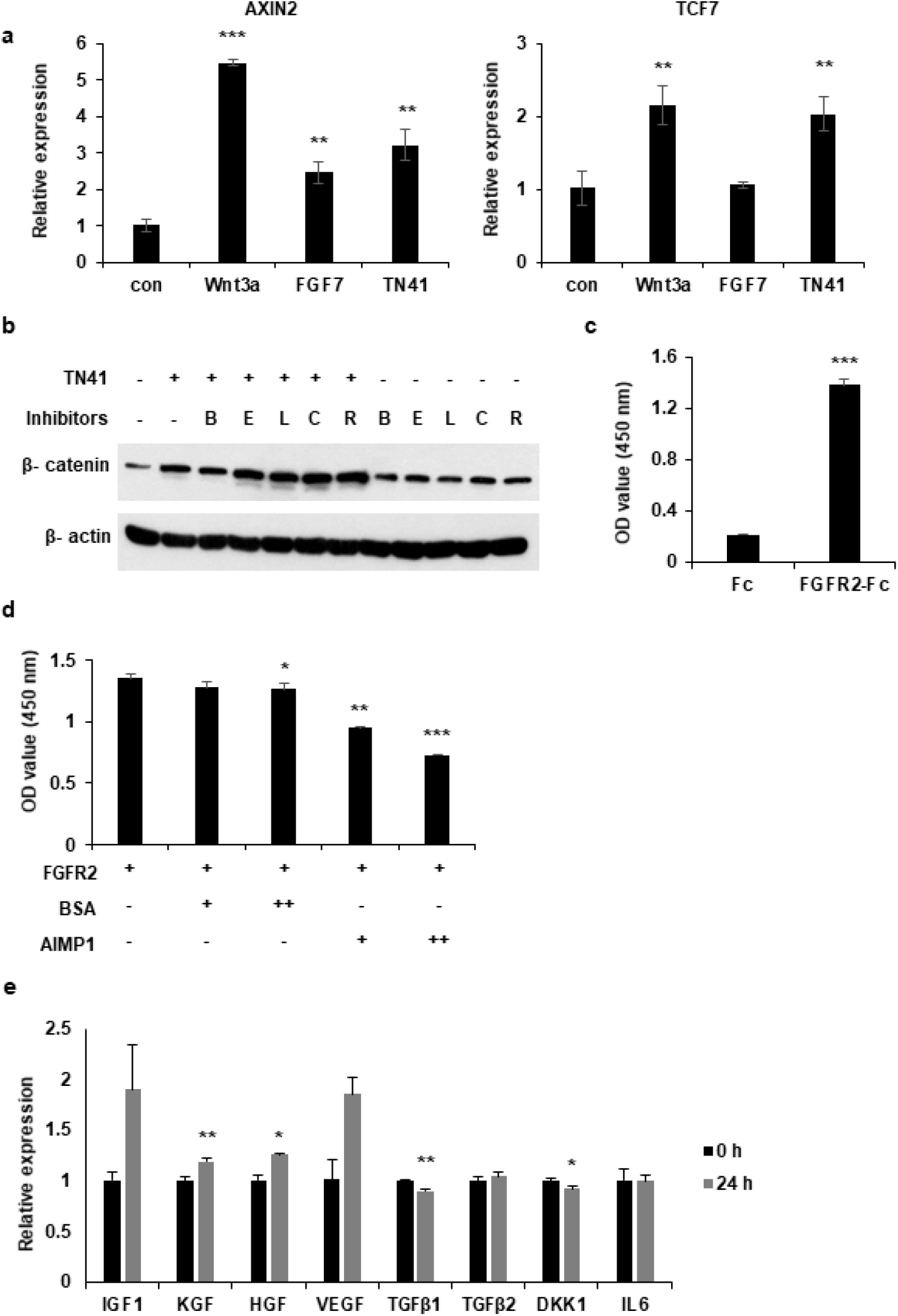
Identification of target cell and receptor of TN41. (**a**) DPCs were treated with 5.1 nM of Wnt3a, 8.9 nM of FGF7, and 20 nM of TN41 for 14 h, and relative gene expression levels were measured by quantitative PCR. (**b**) DPCs were treated with the indicated chemicals (1 μM each), and TN41 (20 nM). B: BGJ398 (FGFR1 and 2 inhibitor), E: Erlotinib (EGFR inhibitor), L: LDN-193189 (ALK2 and 3 inhibitor), C: Crenolanib (PDGFRa and b inhibitor), R: Repsox (ALK5 inhibitor). WB was performed with an anti-beta catenin antibody. (**c**) Interaction of AIMP1 with FGFR2 was determined by ELISA. Biotinylated AIMP1 was added into Fc and FGFR2-Fc coated plates. Interaction of AIMP1 with FGFR2 was detected by a streptavidin HRP antibody. (**d**) Attenuation of interaction of FGF7 with FGFR2 by AIMP1was measured by ELISA. FGFR2-Fc was added into FGF coated plates. Interaction of FGF7 with FGFR2 was detected by anti-human Fc HRP antibody. The effect of AIMP1 on the FGF7 and FGFR2 interaction was determined by adding different amount of AIMP1 (+: 1 μM and ++: 10 μM). (**e**) 20 nM of TN41 was treated into DPCs for 24 h. Cells were harvested and RNA was isolated from cells. Isolated RNA was used as template for synthesizing cDNA. cDNA was amplified with genes which secreted from DPC. Error bars indicate mean +/- SD. ***: P < 0.001, **: P < 0.01, *: P < 0.05.

## REFERENCES

1. Oh JW, et al. A Guide to Studying Human Hair Follicle Cycling In Vivo. J Invest Dermatol 136, 34–44 (2016).

2. Obara K, et al. Hair-follicle-associated pluripotent stem cells derived from cryopreserved intact human hair follicles sustain multilineage differentiation potential. Sci Rep 9, 9326 (2019).

3. Hsu YC, Li L, Fuchs E. Emerging interactions between skin stem cells and their niches. Nat Med 20, 847–856 (2014).

4. Giangreco A, Qin M, Pintar JE, Watt FM. Epidermal stem cells are retained in vivo throughout skin aging. Aging Cell 7, 250–259 (2008).

5. Ji J, Ho BS, Qian G, Xie XM, Bigliardi PL, Bigliardi-Qi M. Aging in hair follicle stem cells and niche microenvironment. J Dermatol 44, 1097–1104 (2017).

6. Turner GA, Bhogal RK. Hair and Aging. Skinmed 14, 338–343 (2016).

7. Jansson L, Kim GS, Cheng AG. Making sense of Wnt signaling-linking hair cell regeneration to development. Front Cell Neurosci 9, 66 (2015).

8. Botchkarev VA, Kishimoto J. Molecular control of epithelial-mesenchymal interactions during hair follicle cycling. J Investig Dermatol Symp Proc 8, 46–55 (2003).

9. Harshuk-Shabso S, Dressler H, Niehrs C, Aamar E, Enshell-Seijffers D. Fgf and Wnt signaling interaction in the mesenchymal niche regulates the murine hair cycle clock. Nat Commun 11, 5114 (2020).

10. Quevillon S, Agou F, Robinson JC, Mirande M. The p43 component of the mammalian multi-synthetase complex is likely to be the precursor of the endothelial monocyte-activating polypeptide II cytokine. J Biol Chem 272, 32573–32579 (1997).

11. Matschurat S, et al. Regulation of EMAP II by hypoxia. Am J Pathol 162, 93–103 (2003).

12. Kim E, Kim SH, Kim S, Cho D, Kim TS. AIMP1/p43 protein induces the maturation of bone marrow-derived dendritic cells with T helper type 1-polarizing ability. J Immunol 180, 2894–2902 (2008).

13. Kim E, Kim SH, Kim S, Kim TS. The novel cytokine p43 induces IL-12 production in macrophages via NF-kappaB activation, leading to enhanced IFN-gamma production in CD4+ T cells. J Immunol 176, 256–264 (2006).

14. Park SG, et al. Dose-dependent biphasic activity of tRNA synthetase-associating factor, p43, in angiogenesis. J Biol Chem 277, 45243–45248 (2002).

15. Park SG, et al. Hormonal activity of AIMP1/p43 for glucose homeostasis. Proc Natl Acad Sci U S A 103, 14913–14918 (2006).

16. Iqbal Z, et al. Missense variants in AIMP1 gene are implicated in autosomal recessive intellectual disability without neurodegeneration. Eur J Hum Genet 24, 392–399 (2016).

17. BoAli A, Tlili-Graiess K, AlHashem A, AlShahwan S, Zuccoli G, Tabarki B. Novel Homozygous Mutation of the AIMP1 Gene: A Milder Neuroimaging Phenotype With Preservation of the Deep White Matter. Pediatr Neurol 91, 57–61 (2019).

18. Park SG, et al. The novel cytokine p43 stimulates dermal fibroblast proliferation and wound repair. Am J Pathol 166, 387–398 (2005).

19. Han JM, Park SG, Lee Y, Kim S. Structural separation of different extracellular activities in aminoacyl-tRNA synthetase-interacting multi-functional protein, p43/AIMP1. Biochem Biophys Res Commun 342, 113–118 (2006).

20. Kim SY, et al. ARS-interacting multi-functional protein 1 induces proliferation of human bone marrow-derived mesenchymal stem cells by accumulation of beta-catenin via fibroblast growth factor receptor 2-mediated activation of Akt. Stem Cells Dev 22, 2630–2640 (2013).

21. Joost S, et al. The Molecular Anatomy of Mouse Skin during Hair Growth and Rest. Cell Stem Cell 26, 441–457 e447 (2020).

22. Schwarz MA, Lee DD, Bartlett S. Aminoacyl tRNA synthetase complex interacting multifunctional protein 1 simultaneously binds Glutamyl-Prolyl-tRNA synthetase and scaffold protein aminoacyl tRNA synthetase complex interacting multifunctional protein 3 of the multi-tRNA synthetase complex. Int J Biochem Cell Biol 99, 197–202 (2018).

23. Muller-Rover S, et al. A comprehensive guide for the accurate classification of murine hair follicles in distinct hair cycle stages. J Invest Dermatol 117, 3–15 (2001).

24. Wang X, et al. Macrophages induce AKT/beta-catenin-dependent Lgr5(+) stem cell activation and hair follicle regeneration through TNF. Nat Commun 8, 14091 (2017).

25. Park SG, Choi EC, Kim S. Aminoacyl-tRNA synthetase-interacting multifunctional proteins (AIMPs): a triad for cellular homeostasis. IUBMB Life 62, 296–302 (2010).

26. Won CH, et al. Comparative secretome analysis of human follicular dermal papilla cells and fibroblasts using shotgun proteomics. BMB Rep 45, 253–258 (2012).

27. Schilling O, Overall CM. Proteome-derived, database-searchable peptide libraries for identifying protease cleavage sites. Nat Biotechnol 26, 685–694 (2008).

28. Zhao P, Alam MB, Lee SH. Protection of UVB-Induced Photoaging by Fuzhuan-Brick Tea Aqueous Extract via MAPKs/Nrf2-Mediated Down-Regulation of MMP-1. Nutrients 11, (2018).

29. Limb GA, et al. Matrix metalloproteinase-1 associates with intracellular organelles and confers resistance to lamin A/C degradation during apoptosis. Am J Pathol 166, 1555–1563 (2005).

30. Bassiouni W, Ali MAM, Schulz R. Multifunctional intracellular matrix metalloproteinases: implications in disease. FEBS J, (2021).

31. Li YH, Zhang K, Ye JX, Lian XH, Yang T. Wnt10b promotes growth of hair follicles via a canonical Wnt signalling pathway. Clin Exp Dermatol 36, 534–540 (2011).

32. Teh MT, et al. Role for WNT16B in human epidermal keratinocyte proliferation and differentiation. J Cell Sci 120, 330–339 (2007).

33. Guo H, Xing Y, Deng F, Yang K, Li Y. Secreted Frizzled-related protein 4 inhibits the regeneration of hair follicles. PeerJ 6, e6153 (2019).

34. Nakamura RE, Hackam AS. Analysis of Dickkopf3 interactions with Wnt signaling receptors. Growth Factors 28, 232–242 (2010).

35. Greco V, et al. A two-step mechanism for stem cell activation during hair regeneration. Cell Stem Cell 4, 155–169 (2009).

36. Suzuki K, et al. Defective terminal differentiation and hypoplasia of the epidermis in mice lacking the Fgf10 gene. FEBS Lett 481, 53–56 (2000).

37. Qu Y, et al. The dual delivery of KGF and bFGF by collagen membrane to promote skin wound healing. J Tissue Eng Regen Med 12, 1508–1518 (2018).

38. Kawano M, et al. Comprehensive analysis of FGF and FGFR expression in skin: FGF18 is highly expressed in hair follicles and capable of inducing anagen from telogen stage hair follicles. J Invest Dermatol 124, 877–885 (2005).

39. Ota Y, Saitoh Y, Suzuki S, Ozawa K, Kawano M, Imamura T. Fibroblast growth factor 5 inhibits hair growth by blocking dermal papilla cell activation. Biochem Biophys Res Commun 290, 169–176 (2002).

40. Janich P, et al. Human epidermal stem cell function is regulated by circadian oscillations. Cell Stem Cell 13, 745–753 (2013).

41. McGee HM, et al. IL-22 promotes fibroblast-mediated wound repair in the skin. J Invest Dermatol 133, 1321–1329 (2013).

42. Fu Y, et al. Structure of the ArgRS-GlnRS-AIMP1 complex and its implications for mammalian translation. Proc Natl Acad Sci U S A 111, 15084–15089 (2014).

43. Philpott MP, Sanders D, Westgate GE, Kealey T. Human hair growth in vitro: a model for the study of hair follicle biology. J Dermatol Sci 7 Suppl, S55–72 (1994).

44. Magerl M, Kauser S, Paus R, Tobin DJ. Simple and rapid method to isolate and culture follicular papillae from human scalp hair follicles. Exp Dermatol 11, 381–385 (2002).

45. Kwack MH, et al. Dihydrotestosterone-inducible dickkopf 1 from balding dermal papilla cells causes apoptosis in follicular keratinocytes. J Invest Dermatol 128, 262–269 (2008).

